# Novel devaluation methods to explore habits in humans

**DOI:** 10.64898/2026.01.25.701564

**Authors:** Mario Michiels

## Abstract

Habits in humans are commonly studied through outcome devaluation paradigms, but most existing tasks fail to capture the robustness of habitual behavior seen in animal models. I introduce two novel behavioral tasks designed to overcome these limitations. In the first task, ("shooting aliens task", n = 45), I simplified an existing instrumental learning task and implemented a novel intra-block reversal method in which stimulus positions changed unexpectedly within blocks while maintaining the same stimulus–action mappings. Participants also completed a classical devaluation phase with explicit reward changes. In the second task ("hands-attack task", n = 44), which relied on real-life avoidance behavior, devaluation was achieved by reversing reward contingencies and allowing participants to inhibit the dominant avoidance response in favor of a more effortful counterattack. Across both tasks, overtrained conditions led to more errors and longer response times after devaluation, confirming increased insensitivity to outcome change. Intra-block reversals in the shooting aliens task produced stronger habitual signatures than standard whole-block devaluation, revealing a greater cost of overriding automatic responses. In the hands-attack task, even without prior training, participants showed clear markers of habitual behavior, suggesting that real-world action patterns can replicate key features of laboratory habits. Interestingly, participants were more accurate in overriding overtrained responses when attacks were highly familiar, possibly due to enhanced perceptual processing, although this came at the cost of longer response times. These findings introduce two complementary tools that address key limitations in current paradigms: the intra-block reversal increases habit sensitivity without inflating working memory demands, while the hands-attack task captures naturalistic habit expression without artificial training, using a single, ecologically valid session. Both are suited for clinical applications, particularly where time constraints or cognitive load limit the feasibility of traditional approaches.

## Introduction

We often need to change our habits (e.g., taking a different route to work due to road construction. Initially, previously learned motor patterns interfere with the new route, reflecting the persistence of the old habit. The additional cognitive cost required to suppress established habits underlies the conflict of exploiting two distinct competing systems (Bernard W. Balleine and O’Doherty 2010). One is the goal-directed system, whereby actions are executed by prospectively evaluating the behavioral options to achieve task goals, considering the available knowledge of the new route and traffic conditions. Goal-directed actions are therefore computationally expensive and slow. In contrast, habits like our usual commute are learned gradually by repetition, are computationally efficient and therefore very fast to execute (Cushman and Morris 2015). Yet, they come at a cost of being highly inflexible and difficult to revert (Seger and Spiering 2011), making it challenging to quickly adjust to the new route.

However, despite extensive research, current human paradigms rarely capture the robustness of habitual behavior observed in animal studies (de Wit et al. 2018; Pool et al. 2022), indicating a need for improved experimental approaches. Here, I introduce two novel behavioral tasks designed to overcome this limitation: a simplified instrumental learning task with novel intra-block reversals (“shooting aliens task”) and a single-session task based on real-life avoidance (“hands attack task”).

The gold standard for studying habits in animals has always been the outcome devaluation procedure. When animals start learning new stimulus–response–outcome (S–R–O) associations (goal-directed behavior), extended training gradually brings behavior under habitual rather than goal-directed control. Then, the outcome is devalued, either by satiety or aversion, and if responding persists, it shows insensitivity to the outcome value, a hallmark of habitual behavior (Adams 1982; B. W. Balleine and Dickinson 1998).

While several parallelisms exist on how humans create everyday life habits, it has been proven difficult to induce habits in humans in laboratory settings using similar animal protocols (de Wit et al. 2018; Pool et al. 2022). Humans are much more likely to change their behavior strategically when reward contingencies change (Seger and Spiering 2011). Alternative approaches to study habit formation as a function of amount of training have been proposed, including action switching and response inhibition tasks (de Wit et al. 2018). Yet, most of the tested protocols did not show the pattern of the expected with overtraining (de Wit et al. 2018; Pool et al. 2022; Poppy Watson 2024): insensitivity to devalued outcomes and therefore an increased difficulty when changing the habitual response.

The validity of previous studies is therefore compromised, as it is unclear whether they are measuring habit formation or something else: A valid behavioral marker for the functioning of the habit system should be sensitive to the amount of instrumental training. To date, two promising novel methods are now available to study habits experimentally in humans which have shown the expected transition from goal-directed to habitual control in function of the amount of training. By adding time pressure during a reversal test, the slower goal-directed system does not get sufficient time to control the habitual action. When comparing different response preparation times before different habitual and non-habitual movements are upcoming (Hardwick et al. 2019) results showed increased probability of making an habitual error for certain preparation times (not too short, not too long). If preparation time was too short, participants responded at random regardless of the amount of previous training. Importantly, when participants had some additional time for preparing their responses, they produced the habitual response, and this effect was larger in extended training conditions. Finally, participants could produce the correct goal-directed response more easily with longer preparation times, hence the effect of the amount of training was attenuated in these trials.

Alternatively, Luque et al. 2020 also manipulated the amount of training and tested habit formation in a partial reversal test (in which one outcome was devalued, see Methods for more details). They did not control response preparation time as (Hardwick et al. 2019) did. Instead, they imposed a very short time for responding. Luque et al. (2020) found that participants suffered extra-cost in response times (RT) when switching the habitual responses during a devaluation procedure (RT-Switch cost) (Luque et al. 2020) and, importantly, this RT-Switch cost was larger in overtrained conditions. This effect has been replicated in a pre-registered study by an independent lab (Nebe et al. 2024). I followed this approach and confirmed it can be a valid habitual measure in a neuroimaging study (Michiels et al. 2025). This extra cost in RT suggests increased difficulty when changing our habitual response, which makes it the perfect candidate for assessing the strength of habits.

Analysis of our previous work showed that, while the findings were statistically robust, participants’ feedback revealed potential for improvement. A general problem with tasks studying habits is that we tend to overcomplicate simple tasks to try to induce habitual errors in the devaluation phase (de Wit et al. 2018). This problem arises from the high flexibility humans have for changing our behavior, as previously discussed. However, this overcomplexity comes at a cost: participants may commit a similar number of mistakes for overtrained and standard trained conditions (de Wit et al. 2018), and the errors they made may be due to the complexity of the tasks and not due to habitual errors. To distinguish these two kinds of errors is usually impossible, which ends up introducing noise in our datasets since we consider both kinds of errors as habitual errors. This high flexibility of humans changing behavior also produces a situation where the last trials for each devaluation block are not as unexpected as we would like, since participants begin to learn the new pattern and start forgetting the habit progressively.

For all this, I aimed to modify our existing aliens task from (Michiels et al. 2025), originally based on (Luque et al. 2020), with the following objectives in mind: 1) Simplify the task to only have habitual errors, and no other kinds of errors (i.e. random guesses or not understanding the devaluation procedure). This also comes from the necessity to simplify the task to be used on patients who have cognitive deterioration, such as in Parkinson’s disease (PD) patients, or even in elder subjects whose cognitive level is not as good as the students in their 20s that usually participate in these studies. 2) Introduce a new devaluation method, which must be more unexpected than the classical devaluation. I therefore introduced an intra-block reversal, where in some trials the participants must change their responses, presented in a random manner. From now on, I will refer to this task as the shooting aliens task.

Finally, the way that habitual behavior studies are done often requires multiple days of training. Online tasks facilitate this, but still some patients do not have the necessary means or knowledge to be able to participate in computerized online tasks from their homes. Coming to the hospital multiple consecutive days is also a logistical problem for many patients. Modifying our aliens task to solve this did not seem feasible, since establishing new habitual associations in 30-minute sessions could be challenging and theoretically difficult to explain. Therefore, we would need a task where the associations have been previously learned over our life (i.e., real-life habits: (Guida et al. 2022). To address this, I designed the hands attack task, a computerized adaptation of the classical hands attack game based on lifelong avoidance learning. Because this response is deeply ingrained, no new learning is required. The task also allows the assessment of impulsivity, which is frequently affected in PD patients receiving dopaminergic therapy or surgical treatment (Weintraub et al. 2010).

I hypothesize that mixing devalued and non-devalued trials within the same block (intra-block reversal) will sustain the element of surprise and limit goal-directed compensation, resulting in a higher RT-Switch cost throughout our novel intra-block reversal phase. Moreover, our simplifications in the S-R-O configurations and instructions should reveal an increased number of habitual errors and RT-switch cost, due to minimizing noise from random or comprehension-related errors. Additionally, I expect that these measures (i.e. habitual errors and RT-Switch cost) can be translated to our novel hands attack task, which is based on automatic real-life avoidance behavior. This would show that overlearned real-life habits can be reliably triggered and measured in a controlled lab setting.

The main objective of this study is to develop and validate two novel behavioral tasks that address key limitations in the experimental study of habits in humans. First, to induce more habitual errors and maximize the RT-Switch cost as a cleaner behavioral marker of habit strength by simplifying our instrumental learning task (i.e. aliens task) and introducing an intra-block reversal with sudden unexpected changes. And second, by developing a task based on automatic avoidance responses rooted in real-life habits (i.e. hands attack task), which would offer a fast, training-free method that could be easily integrated into clinical research on disorders involving habitual behavior, such as Parkinson’s disease (Redgrave et al. 2010) and impulsivity (Cyders and Coskunpinar 2011).

## Methods

### Participants

#### Shooting aliens task

45 healthy psychology students were recruited from the Universidad Complutense de Madrid (mean age = 19.2 years, SD 1.1; 36 female). I confirmed that the subjects did not have any neurological disorders by general questionaries, and those who had mild-severe disorders were not included in the study (including those with moderate disorders but in medication).

#### Hands attack task

44 healthy psychology students were recruited from the Universidad Complutense de Madrid (mean age = 19.4 years, SD 1.3; 40 female). I confirmed that the subjects did not have any neurological disorders by general questionaries, and those who had mild-severe disorders were not included in the study (including those with moderate disorders but in medication).

### Task

The shooting aliens task refines the aliens task (Michiels et al. 2025) (originally based on (Luque et al. 2020))) by simplifying its design and introducing novel devaluation methods. In this reward learning task, participants are trained on stimulus-response-outcome associations. For each trial, one stimulus (i.e. an alien image) appeared at the center of the screen. At each side of the screen, a weapon was displayed. Participants’ responses consisted of pressing the <q> or <p> buttons (i.e. left/right), thus earning points by shooting the aliens. For each trial, there were only two possible responses (i.e. <q> or <p>). One of them led to killing the aliens (optimal outcome; 100 points), while the other only hurt them (suboptimal outcome; 5 points) (Figure 1A; Table 1). These associations were learned by trial and error. Four alien stimuli were used, distinguishable only by their color (brown, blue, purple, and yellow).

**Figure 1.**
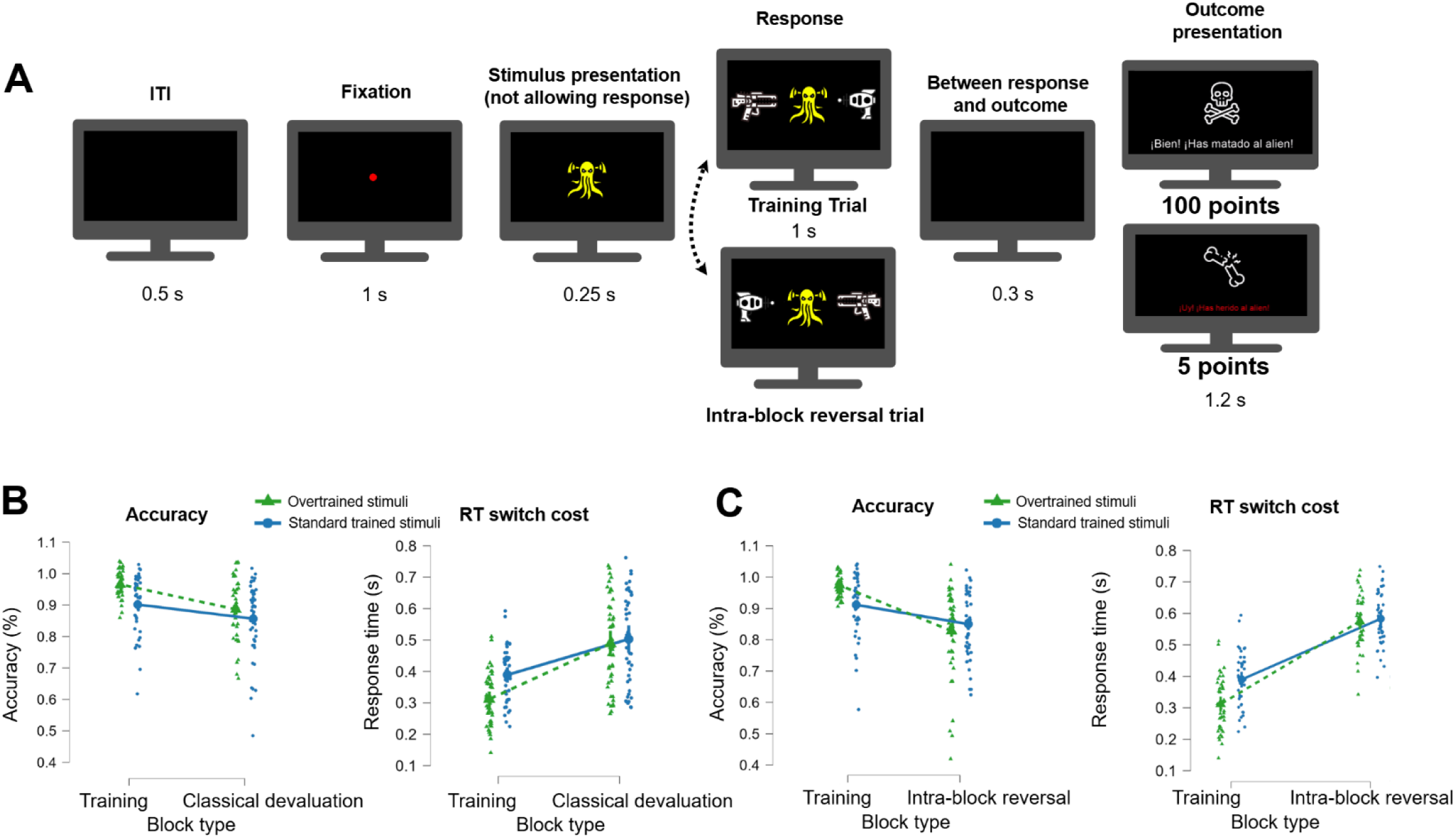
Shooting aliens task and behavioral results. **A)** Example of a single trial. On each trial, an alien appeared at the center of the screen, and two weapons were displayed on the sides. Participants were instructed to press either the <q> (left) or <p> (right) button to choose which weapon to attack with. Choosing the weapon that is deadly to that alien yielded an optimal outcome (100 points); the alternative response only hurts the alien and yielded a suboptimal outcome (5 points). **B)** Comparison between overtrained and standard trained stimuli performance in terms of accuracy and RT for classical devaluation block and **C)** Intra-block reversal. *: Intra-block reversal trial where weapons position was exchanged but participants still had to choose the one that kills the alien. Note: Data points are vertically jittered for visualization purposes; values exceeding 1 on the accuracy axis do not represent actual data above 100% accuracy.

**Table 1.** S-R-O associations of the “shooting aliens” task. S1-S4 refers to the stimulus shown in the center of the screen (i.e. the colored alien image). R1 and R2 refers to selecting the left and right weapon, respectively. For example, S1→R1→O1 means that a red alien has been killed by the weapon on the left side, which delivers the high-value outcome (100 points). Note this example is hypothetical, as all conditions were counterbalanced across participants (see Methods section).

| Stimulus->Response->Points (outcome) | Relative frequency in training | Relative frequency in classical devaluation |
| --- | --- | --- |
| S1->R1->100 (kill) | ×1 (standard trained) | ×1 |
| S1->R2->5 (hurt) |  |  |
| S2->R2->100 (kill) | ×3.5 (overtrained) | ×1 |
| S2->R1->5 (hurt) |  |  |
| S3->R1->100 (kill) | ×1 (standard trained) | ×1 |
| S3->R2->5 (hurt) |  |  |
| S4->R2->100 (kill) | ×3.5 (overtrained) | ×1 |
| S4->R1->5 (hurt) |  |  |

Following the same setup as in (Luque et al. 2020) and (Michiels et al. 2025), stimuli were divided into two training conditions: overtrained and standard trained. Overtrained stimuli appeared 3.5 times more frequently than standard ones (Table 1), and their assignment was counterbalanced across participants.

Participants completed 3 online sessions in consecutive days (∼ 30 minutes each), where each of the first two consisted of 4 training blocks. The last session consisted of 3 training blocks (+1 one between devaluation blocks), 3 classical devaluation blocks and 3 intra-blocks reversals (Table 2). Each block contained 44 trials. To maintain engagement, participants were given bonus points after each block if their accuracy exceeded 70% and their mean reaction time was below 0.45 s per trial. In addition, the three highest-scoring participants received a monetary reward of 25 € at the end of the final session.

**Table 2.** Blocks structure for the last session of the “shooting aliens” task.

| Number of block | Type |
| --- | --- |
| 1 | Training |
| 2 | Training |
| 3 | Intra-block reversal |
| 4 | Training |
| 5 | Intra-block reversal |
| 6 | Training |
| 7 | Classical devaluation (high value outcome) |
| 8 | Training |

For the two first sessions, I included a very short devaluation block (3 trials) to make participants aware that they must learn the S-R-O associations instead of just S-R associations (i.e. I try to avoid that they simply learn left-right associations for each stimulus). This short devaluation block has no other real use and does not train participants for the real devaluation procedure on the third day. The third session was the only one that included the actual devaluation procedures.

In this last session, participants first completed two training blocks, to remind them of the already learned associations. Then, we introduced our novel intra-block reversal, where some weapon images randomly change their side (Figure 1A). This is an unexplicit devaluation method, which is done on a per-trial basis instead of a whole block. 50 % of trials in these blocks present the stimuli exchanging their position, but the order of these is randomized. These two new features (i.e. inexplicitly and randomly) since participants are not expecting any sudden changes, and they will not be able to re-learn the new associations.

Finally, we presented a classical devaluation block (see Table 2), where a slide appeared on the screen instructing participants that one of the optimal outcomes (i.e., killing the aliens) was now forbidden by the police and would yield zero points. Importantly, this phase revealed the purpose of having both overtrained and standard trained conditions, showing how the amount of training affects behavior during devaluation. Moreover, in this phase, we balanced the number of overtrained and standard trained trials to improve discriminability be-tween conditions.

#### Hands attack task

The hands attack task is a videogame version of the popular hands attack game (also called hands warmer), with the only difference that here the participants only play the role of the defense side. The task is designed for both healthy subjects and patients since it is simple and does not require a high memory workload. Only one online session was required (∼ 30 minutes), which consisted of 10 blocks, each with 36 trials (Table 3).

**Table 3.**
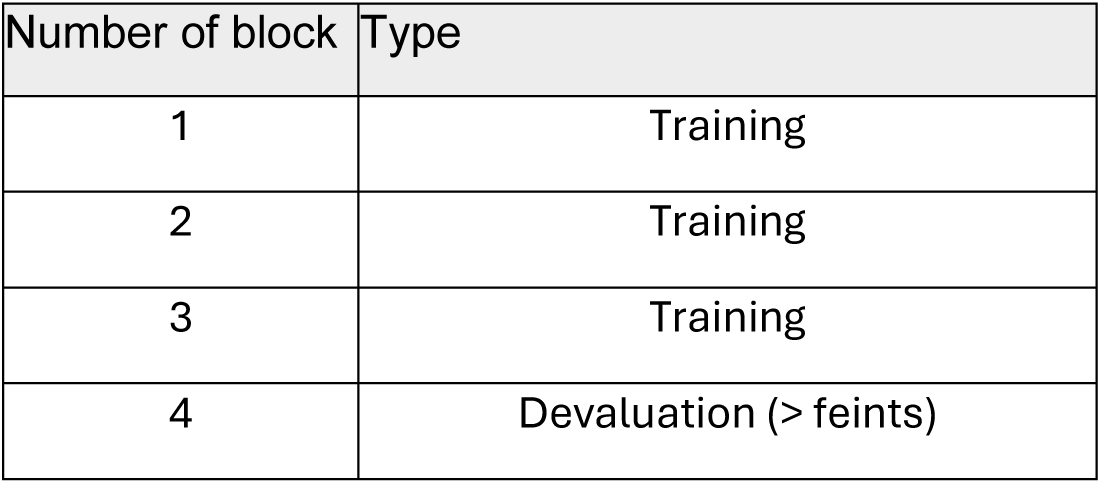
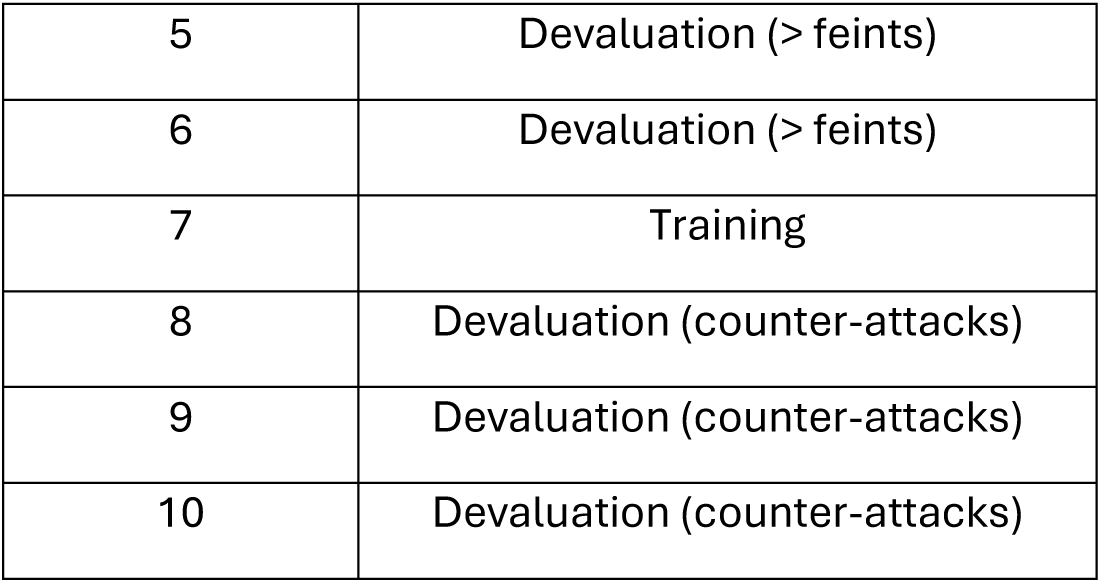
Blocks structure for the “hands attack” task.

Each trial commences once we have both <q> and <p> keys pressed (Figure 2A). Keeping the keys pressed simulate that your hands are still on the screen. The opponent hands are shown on the upper half side of the screen while the participant hands are in the lower half of the screen. Whenever one of the opponent hands move towards the participant hands, participants must withdraw the attacked hand by pressing the left/right button to dodge the attack.

**Figure 2.**
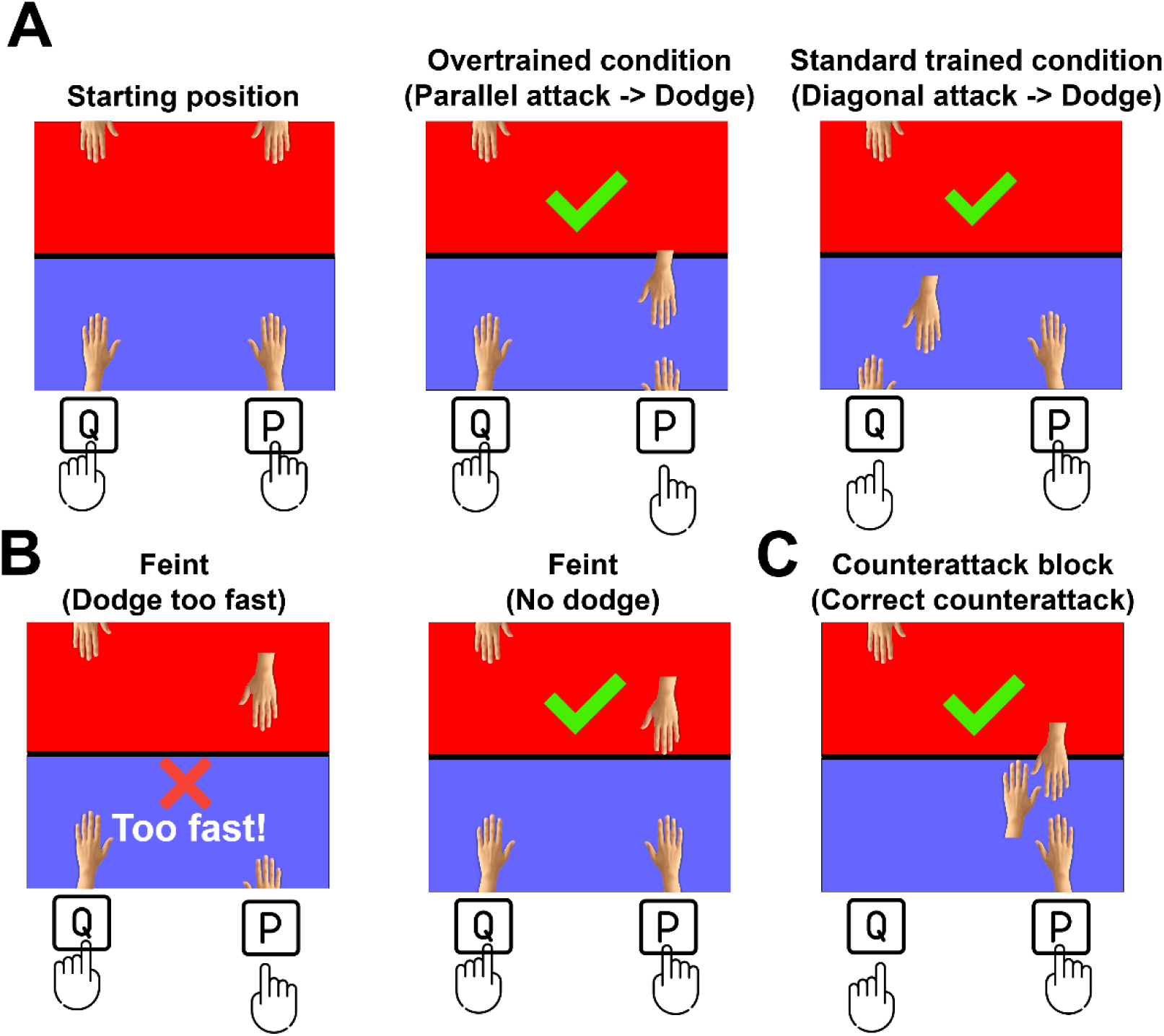
Hands attack task. Single trials examples**. A)** All trials commence by keeping both keys pressed and in the training phase they must dodge parallel and diagonal attacks. Participants also must take care to only respond whenever the attacker hand has surpassed the middle line since 30 % of trials are feint trials that fake real attacks but actually stop in the middle line. for **B)** Devaluation blocks where feint trials were increased to 70 % **C)** Trial where participants were allowed to counter-attack if they wanted a higher reward than just dodging, which was still allowed.

To include overtrained and standard trained stimuli in this task, I had to find a more creative way of doing so. Hence, I implemented parallel attacks as overtrained stimuli and diagonal attacks as the standard trained (Figure 2A), with parallel attacks appearing 3.5 times more than diagonals. Both parallel and diagonal attacks are performed at the same speed. If your hand was not withdrawn in less than 0.5 s, the attacked hand will receive the impact, and no points will be obtained. Withdrawing both hands invalidates the trials and yields no points.

However, there is a time window (∼ 0.25 s) where participants must not withdraw the hand too early, as some trials stop in the middle line and retreat to the starting position (i.e. feints) (Figure 2B). For these trials, the reward is obtained just by not releasing any keys (i.e. not withdrawing the hands) (Figure 2B).

The task starts with the training phase (Table 3), which consists of 3 blocks where 30 % of trials per block are feint trials while the rest are real attacks. The order of the feint trials is always randomized. Then, three devaluation blocks are shown, where the percentage of feint trials is exchanged with the real attacks (i.e. 70 % of the trials are feints). A normal training block is presented after this, so participants can have a clearer state before the final devaluation. This final devaluation consists of 3 training blocks where participants are now allowed to counter-attack using the opposite hand to the one that is being attacked. Dodging the attack by just withdrawing the targeted hand is always an option, as in the rest of the experiment. However, a correct counter-attack will yield 10 times more points than dodging the attack. Note that this involves an additional risk since the counter-attacks require more distance and coordination to hit the opposite hand.

The points for each correct trial are calculated also by considering the RT, with the following formula: *base_points - (response_time*4)*. A monetary reward of 25 € was given to each of the top 3 participants in terms of points in the last session.

### Behavioral analysis

A linear mixed effect model (LMM) was used to test the statistical significance of the behavioral effects across the two tasks. I selected this model as other forms of comparisons (e.g. ANOVAs) are not ideal to test our hypothesis given the current design does not include even number of trials across conditions (i.e. higher trial amount in overtrained than standard trained stimuli).

#### Shooting aliens task

For the shooting aliens task, the focus lies in the analysis of the third-day session, where participants had already established the associations and the devaluations occurred. The dependent variable was response accuracy and response time (RT) in 2 separate models. The fixed effect variables were stimulus type (overtrained or standard trained) and block type (training or devaluation) with a random effect grouping factor as the subject ID. At the same time, this analysis was run twice, first for classical devaluation blocks and then for our novel intra-block reversal. Additionally, the first block was excluded from the analysis as participants were still learning the task and would only introduce undesired noise in the analysis.

For the response accuracy analysis, I initially attempted to fit a maximal random effects structure, including random slopes for block_type, overtraining_stim, and their interaction, as well as random intercepts for participants. However, this complex model structure led to a singular model fit, indicating that the specified random effects structure was too complex for the available data. To address this, I simplified the random effect’s structure by removing the random slope for the interaction. The final model is as follows, for each devaluation type:

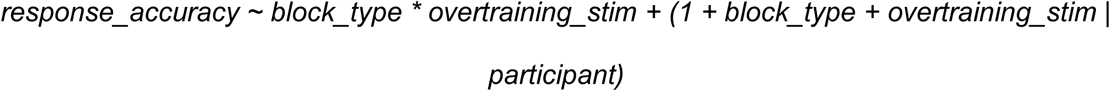

As for the response time analysis, the model is as follows, for each devaluation type:

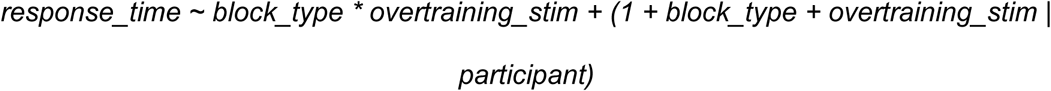

The result from this model is what I refer to as the RT switch cost.

#### Hands attack task

The analysis of the hands attack task focused on three conditions of particular interest. First, it is crucial to analyze the devaluation blocks where feint trials were increased to 70 % instead of the usual 30 %. Here, the first block was excluded from the analysis as participants were still learning the task and would only introduce undesired noise in the analysis. Similarly, I excluded the 6^th^ block as it was just a training block in the middle of both devaluation types, whose purpose was just to remind the subjects the original state of the task. To avoid the singular model fit issue, I included random slopes for block_type, overtraining_stim, but not their interaction. The final model is for the accuracy is as follows:

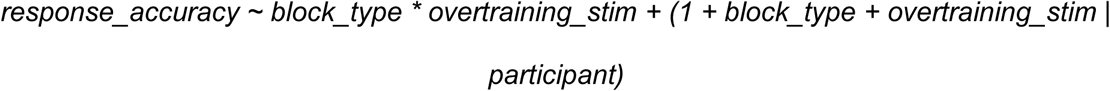

As for the response time analysis, the model is as follows, for each devaluation type:

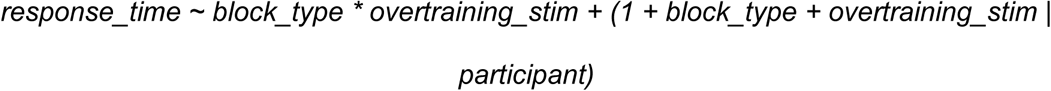

Then, the primary focus of interest was directed toward the blocks featuring counterattacks. For this, I first analyzed the trials where participants tried to counterattack. To avoid the singular model fit issue, I only included random intercepts for participants. The final model is for the accuracy is as follows:

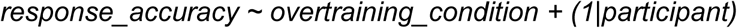

As for the response time:

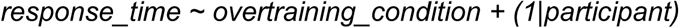

Finally, I wanted to investigate the trials from the counterattacks blocks where participants tried to dodge instead of counter-attacking. To have a fair comparison in terms of accuracy, I excluded the feint trials from the training blocks, since there are no feints in the counter-attack blocks. To avoid the singular model fit issue, I included random slopes for block_type, overtraining_stim, but not their interaction. The final model is for the accuracy is as follows:

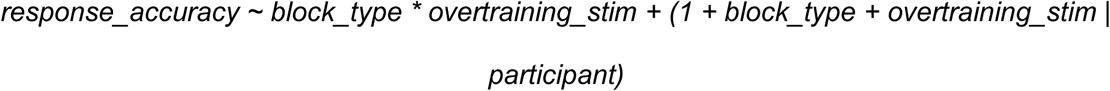

As for the response time analysis, the model is as follows, for each devaluation type:

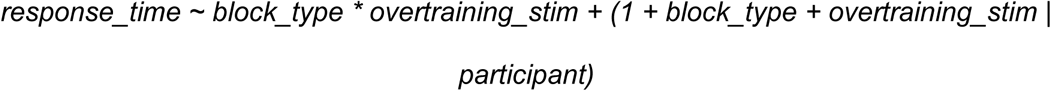

## Results

### Shooting aliens task

First, we verified that participants successfully learned the associations during the training phase. This was revealed by a higher accuracy for overtrained stimuli with respect to the standard trained (Estimate = 0.061, SE = 0.009, z = 6.642, p < .001). This good learning performance in the training phase was confirmed by lower RTs for the correct trials in the overtrained condition (Estimate = -0.105, SE = 0.007, z = -15.916, p < .001). This is in line with the expected results for well-trained associations (i.e. precise and fast responses).

For the accuracy in the classical devaluation blocks, participants committed more mistakes for the overtrained stimuli, with respect to their accuracy in the training phase (i.e. less response switches), as shown by an interaction between block x amount of training (F(1, 9764.03) = 8.83, p = 0.003; Figure 1B). That is, changing the habitual answer was indeed more difficult for the overtrained condition, which is in line with our hypothesis. This effect was more intense for our novel intra-block reversal (F(1, 9856.11) = 48.32, p < 0.001; Figure 1C), which is in line with our hypothesis that unexpected changes will induce more habitual errors.

For the RT in the classical devaluation blocks, participants had higher RT for the response switches in overtrained stimuli when compared to their accuracy training phase (i.e. RT switch cost), as shown by an interaction between block x amount of training (F(1, 9237.14) = 63.88, p < 0.001; Figure 1B). That is, changing the habitual answer was indeed more difficult in terms of RT for the overtrained condition, which is also in line with our hypothesis. The intensity of the effect is again a bit higher than our novel intra-block reversal (F(1, 9180.59) = 75.92, p < 0.001; Figure 1C). The results align with our hypothesis that there exists more RT switch cost for extended training cues, and an even higher extra cost in time when this habitual change is unexpected.

### Hands attack task

First, we confirmed that learning was properly established during the training phase. This was revealed by a higher accuracy for overtrained stimuli (parallel attacks) with respect to the standard trained (diagonal attacks) (Estimate = 0.148, SE = 0.019, z = 7.853, p < .001). This is in line with the expected results for optimal learning (i.e. precise responses). Further exploring the learning dynamics, I observed that RTs were similar for the overtrained and standard trained conditions when effectively dodging an attack (Estimate = -0.105, SE = 0.007, z = -0.408, p =0.683). This may be due to the intrinsic nature of the task where the reactions were somewhat impulsive as reacting to a danger, a behavior we have already learned during all our life.

For the devaluation blocks where the number of feints were significantly increased, our main interest was in analyzing the number of correct dodges on trials without feints. For this, participants had more mistakes for the overtrained stimuli (parallel attacks), with respect to their accuracy in these stimuli in the training phase (i.e. less response switches), as shown by the interaction block x amount of training (F(1, 3527.22) = 53.7, p < 0.001; Figure 3A). Moreover, the correct dodges in these blocks required more time than in training phase, but only for the overtrained condition, as revealed by the interaction block x amount of training (F(1, 3046.31) = 10.21, p = 0.001; Figure 3A). That is, changing the habitual answer was indeed more difficult for the overtrained condition, which is in line with our hypothesis.

**Figure 3.**
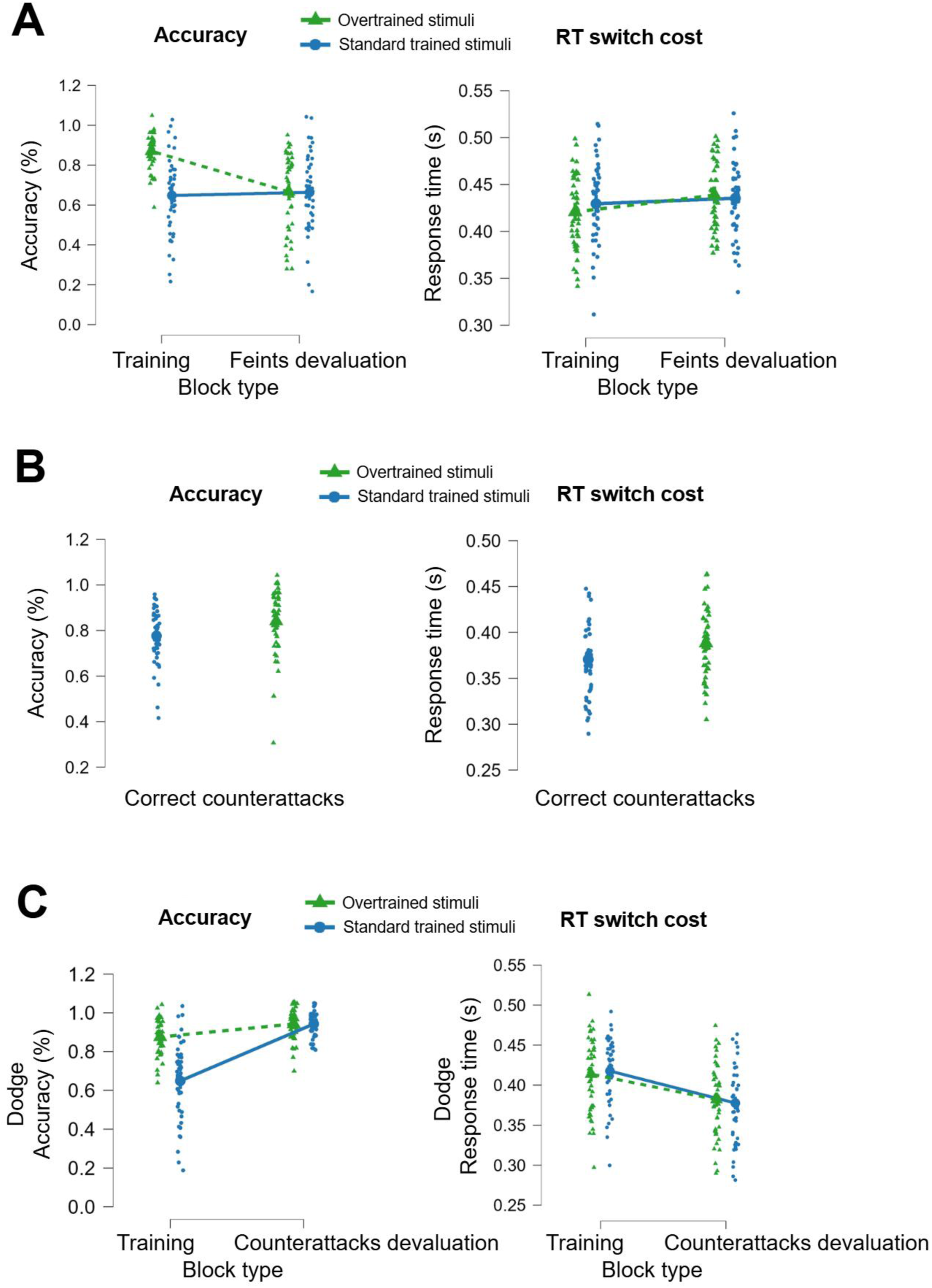
Hands attack’s behavioral results. Comparison between overtrained and standard trained stimuli performance in terms of accuracy and RT for **A)** Devaluation blocks where feint trials were increased to 70 % instead of the usual 30 %; **B)** Trials where participants performed a correct counterattack, from the blocks where they were allowed to do so; **C)** Trials from the counterattacks blocks where participants tried to dodge instead of counter-attacking. Note: Data points are vertically jittered for visualization purposes; values exceeding 1 on the accuracy axis do not represent actual data above 100% accuracy.

For the blocks in which counterattacks were enabled by pressing the opposite button, the first step was to analyze the number of correct counterattacks. I found a higher number of correct counter-attacks for the overtrained condition (parallel attacks) with respect to the standard trained condition (diagonal attacks), as shown by the amount of training condition (F(1, 2185.4) = 16.26, p < 0.001; Figure 3A). However, the response time was higher when counter-attacking in the overtrained condition (F(1, 2168.09) = 73.35, p < 0.001; Figure 3B). This is in line with the theory of requiring an extra cost in time to change our habitual answer. Finally, when no counter-attack was tried, we can see that the accuracy for dodging the attacks in the standard trained condition was much higher than in the training blocks, as revealed by the interaction block x amount of training (F(1, 4581.66) = 123.52, p < 0.001; Figure 3C). This may seem contradictory at first glance, but a possible explanation is the already good performance of dodging parallel attacks (i.e. overtrained condition) where there was almost no room to improve. Also note that in these blocks were counter-attacks were allowed, there were no feints, so the overall accuracy for dodging the attacks is always higher. The response times for dodging the attacks in these blocks is lower, as expected, since no feints were present. However, this difference was more intense for the standard trained condition, as shown by the interaction block x amount of training (F(1, 3954.29) = 5.78, p < 0.016; Figure 3C). This may be again because the RTs for standard trained condition (i.e. dodging diagonal attacks) had greater potential for improvement once they were no feints-

## Discussion

I developed two novel tasks to test innovative approaches to overcome the challenges in translating outcome devaluation experiments from animal models to human studies. I revealed that the simplification of an existing associative task (i.e. aliens task) can indeed facilitate more habitual behavior without hurting our goal-directed actions when needed. That is, participants now are more resistant to changing their habitual actions, and even when they succeed, the required extra time to do so is higher (i.e. higher insensitivity to devaluation, both in terms of accuracy and RT). Moreover, the inclusion of intra-block reversals proved that this insensitivity to devaluation is indeed higher than for classical whole-block devaluations, both in terms of higher habitual errors and higher RT-switch cost.

In pursuit of making these advancements easier to apply in clinical settings, I tested whether a novel single session task (i.e. hands attack task) would also align with classical habitual behavior theory. To follow classic habits theory, there would be only two ways of doing so: 1) A tremendously extensive single session where participants learn a new task; 2) A task that resembles a real-world environment for what we already have “trained” through life. The first option would have so many drawbacks that make it suboptimal (e.g. fatigue in the last phases: devaluation, suboptimal memory consolidation since there were no sleeping periods, etc.). However, I revealed that a “real-world” task (i.e. hands attack task), where we do not need to learn the associations and actions, can reproduce the same habit features as the classical lab developed tasks. It is important to note that even if our behavior in this task is based on real-live avoidance learning throughout our life, I had to introduce an artificial overtrained condition (parallel attacks) to compare it with a standard trained condition (diagonal attacks). So, even if no training was required to learn the associations (i.e. withdraw one’s hand from danger), perceiving that there is an overtrained condition may be seen as a kind of training. Similarly, we also observe this for the devaluation blocks where I let participants counter-attack using the opposite button.

I hypothesize that learning these conditions is significantly less demanding than learning whole new associations exclusively for a task, and therefore they did not hurt the habitual behavior features. Therefore, the two devaluation tests I implemented (blocks with more feints, blocks allowing counterattacks) show results that align with our hypotheses in terms of insensitivity to devaluation. However, there was the unexpected result of having higher accuracy for counter-attacking (i.e. changing the habitual answer) when the overtrained stimuli were presented (i.e. parallel attacks). A possible explanation for this may be that participants enhanced perceptual processing of parallel attacks made them more confident to perform counter-attacks. These attempts, however, did require an extra cost in time (RT switch cost), which aligns with our hypothesis.

It is now well-known that most previous devaluation studies in humans may not have fully captured the complexity of habitual behavior (Poppy Watson et al. 2022). These concerns may be explained due to their inability to show insensitivity to devaluation in terms of accuracy, RTs, or even both (de Wit et al. 2018; Pool et al. 2022; Gera et al. 2023; P. Watson et al. 2023). We may not call total failures to these attempts, since most did show partial success revealing some habitual features. Lessons from these studies taught us that humans need a task with extensive training and a more challenging devaluation. Extensive training is easily accomplished with multiple days of training but increasing the devaluation difficulty must be carefully done since it may induce participants to behave almost utterly in a goal-directed way. Attempts on doing so incurred in complicating the tasks so much that they actually raised concerns about participants always exhibiting greater goal-directed behavior (de Wit et al. 2018). For this, I introduced our intra-block reversal method, where the unexpected changes fulfill both simplicity and increased challenging conditions. I also ensured that both of our tasks included a clear response cost (i.e., an explicit negative consequence for each response) (Reed 2001; Perez and Dickinson 2020; Pérez and Soto 2020; Pool et al. 2022). A real monetary reward to the top 6 participants in terms of accuracy (with bonus points for a fast total time) helped to ensure this participants’ interest following devaluation.

For this, some authors have opted to deviate from outcome devaluation methods and test habitual behavior with other techniques. These include Go/no-go task with familiar stimuli: (Ceceli, Myers, and Tricomi 2020), motor-sequence learning tasks, maze navigation (Smith and Graybiel 2013), two-stage tasks, contingency degradation, and Pavlovian conditioning – Pavlovian to instrumental transfer (Hogarth, Lam-Cassettari, and Pacitti 2019). The problem when using these techniques is that the devaluation paradigm is still considered the gold-standard for studying habitual behavior in animals, so direct comparisons with these studies have proven difficult, and controlling an extensive training scenario is often not possible.

For outcome devaluation, a few modifications have proven successful. For instance, (Meier, Staresina, and Schwabe 2022) successfully introduced the stress factor in humans to confirm a shift behavioral control from a goal-directed to a habitual system. (Hardwick et al. 2019) tried different response preparation times before different habitual and non-habitual movements are upcoming. (Glück et al. 2021) tried a modified devaluation paradigm with a dual-task to show how existing habits actively interfere with learning new adaptive behaviors. (P. Watson et al. 2023) attempted a symmetrical outcome-revaluation task to improve the slips-of-action paradigm by having cleaner comparisons between devalued/still valuable conditions, but the result did not reveal insensitivity to devaluation. (Gera, Barak, and Schonberg 2024) trained participants on an action sequence rather than a single action, which may contribute more to habit formation as shown by previous research (Dezfouli, Lingawi, and Balleine 2014).

There is also a relatively new trend that aims to make use of tasks that mimic real-world scenarios so no training is required, and therefore, more natural habitual behaviors may be seen. Even when these studies succeed (Gera, Barak, and Schonberg 2024), there was no way to perform a direct comparison with classical devaluation methods. With our hands-attack task I expect to bridge the gap between these two worlds. The inclusion of different conditions (overtrained stimuli, feints, etc.) requires participants to make rapid decisions based on an already learned action through life (i.e. withdraw one’s hand from danger), more closely mimicking real-world habit engagement and disruption (Dolan and Dayan 2013). For our shooting aliens task, the inclusion of our intra-block reversal also mimics real-world scenarios where habitual actions are disrupted by unexpected changes, providing a more naturalistic measure of habit strength. This may also minimize the confounding effects of working memory load present in the slips of action paradigm (Sjoerds et al. 2016) and more accurately assess implicit habit strength (Bernard W. Balleine and O’Doherty 2010).

Importantly, I argue that in the devaluation and intra-block reversal phases, it is not required for overtrained performance/speed to fall below the standard trained condition. In other words, the decline in performance/speed is sufficient to indicate a habit-related cost. The advantage gained through extended practice disappears once contingencies change, revealing reduced flexibility. Statistically, this pattern is best captured by examining the interaction between extended training and contingency change, which our within-subject design maximizes. Only the overtrained condition acquired an advantage, and therefore only this condition shows a selective decline. Without accounting for this interaction, the cost would remain undetected, as it only becomes meaningful relative to baseline performance (i.e. in training blocks). This view is consistent with human studies showing that habit effects are expressed as the disappearance of a prior training-related advantage rather than a drop below standard trained levels (Hardwick et al. 2019; Gera et al. 2023).

Regarding neural circuitries, it would be expected that our intra block devaluation may engage prefrontal cortical regions differently than explicit devaluation procedures. Specifically, it might reveal how the anterior cingulate cortex (ACC) and orbitofrontal cortex (OFC) dynamically update value representations in the absence of explicit cues (Rushworth et al. 2011). This could provide insights into the neural mechanisms underlying the flexibility of seemingly habitual behaviors, a key area of interest in addiction research.

For the hands attack task, the increased motor complexity in our task likely engages a broader network of motor planning and execution regions, including the supplementary motor area (SMA) and premotor cortex (Hardwick et al. 2013). The formation of habits in this task may involve the chunking of action sequences, a process linked to the dorsolateral striatum (Smith and Graybiel 2016).

### Limitations

As noted by other authors, the outcome devaluation method cannot disentangle whether the transition from goal-directed to habitual behavior is due to increased habitual control, decreased goal-directed control, or both (Bernard W. Balleine and Dezfouli 2019; Pool et al. 2022).

I am also aware that the hands-attack task conditions may not be one-to-one equivalent to classical studies with learning and devaluation blocks. To elucidate this, we would have to perform extensive tests with fMRI/EEG to see whether the neural correlates are the same.

### Future research

Applying our tasks to clinical populations, such as those with obsessive-compulsive disorder or addiction, could reveal whether these conditions are characterized by overreliance on habitual control or deficits in goal-directed processing (Gillan et al. 2016; Hogarth, Lam-Cassettari, and Pacitti 2019) used classical outcome devaluation with Pavlovian to Instrumental Transfer to reveal a different theory where addiction would be primarily driven by excessive goal-directed drug choice under negative affect, rather than by habits or compulsion. Our refined tasks could help to shed light on these clinical populations. Critically, our hands attack task would be logistically ideal for clinical settings since it does not involve any additional training, and it only consists of a 30 minutes session. Moreover, this task can also serve for impulsivity assessments, since it provides some characteristics (i.e. feint trials) that can evaluate behavioral control and address the potential relationship between impulsivity and habit formation (Dalley, Everitt, and Robbins 2011).

Finally, future research should explore the role of dopamine in modulating the balance between goal-directed and habitual behavior using our paradigms. Pharmacological manipulations or studies in populations with altered dopamine function (e.g., PD patients) could provide valuable insights into the neurochemical basis of habit formation and flexibility (Deserno et al. 2015). Moreover, since our hands-attack task is purely motor-based, it allows us to assess motor function, even in a lateralized manner. This can be particularly relevant for clinical populations where hemispheric motor control differences are significant, such as in PD (Paparella et al. 2024).

## Author Contributions

The author was solely responsible for all aspects of the work.

## Ethics approval

The study protocol was reviewed and approved by the Comité Ético de Investigación con medicamentos (CEIm) of HM Hospitales (Approval code: CEIm HM Hospitales 22.11.1190E4-GHM; Approval date: 23 November 2022; Meeting Act No. 262; Protocol version 3.0, dated 5 November 2022).

## Consent to participate

All participants provided written informed consent prior to participation. Signed consent forms are securely stored in accordance with institutional and legal requirements.

## Consent for publication

The author gives full consent for publication.

## Availability of data and materials

Data and materials are available upon request.

## Code availability

Code is available upon request.

## Open practices statement

None of the data or materials for the experiments reported here are publicly available, and none of the experiments were preregistered.

## References

Adams, Christopher D. 1982. “Variations in the Sensitivity of Instrumental Responding to Reinforcer Devaluation.” The Quarterly Journal of Experimental Psychology. B, Comparative and Physiological Psychology 34 (2b): 77–98.

Balleine, B. W., and A. Dickinson. 1998. “Goal-Directed Instrumental Action: Contingency and Incentive Learning and Their Cortical Substrates.” Neuropharmacology 37 (4–5): 407–19.

Balleine, Bernard W., and Amir Dezfouli. 2019. “Hierarchical Action Control: Adaptive Collaboration between Actions and Habits.” Frontiers in Psychology 10 (December): 2735.

Balleine, Bernard W., and John P. O’Doherty. 2010. “Human and Rodent Homologies in Action Control: Corticostriatal Determinants of Goal-Directed and Habitual Action.” Neuropsychopharmacology: Official Publication of the American College of Neuropsychopharmacology 35 (1): 48–69.

Ceceli, Ahmet O., Catherine E. Myers, and Elizabeth Tricomi. 2020. “Demonstrating and Disrupting Well-Learned Habits.” PloS One 15 (6): e0234424.

Cushman, Fiery, and Adam Morris. 2015. “Habitual Control of Goal Selection in Humans.” Proceedings of the National Academy of Sciences of the United States of America 112 (45): 13817–22.

Cyders, Melissa A., and Ayca Coskunpinar. 2011. “Measurement of Constructs Using Self-Report and Behavioral Lab Tasks: Is There Overlap in Nomothetic Span and Construct Representation for Impulsivity?” Clinical Psychology Review 31 (6): 965–82.

Dalley, Jeffrey W., Barry J. Everitt, and Trevor W. Robbins. 2011. “Impulsivity, Compulsivity, and Top-down Cognitive Control.” Neuron 69 (4): 680–94.

Deserno, Lorenz, Quentin J. M. Huys, Rebecca Boehme, Ralph Buchert, Hans-Jochen Heinze, Anthony A. Grace, Raymond J. Dolan, Andreas Heinz, and Florian Schlagenhauf. 2015. “Ventral Striatal Dopamine Reflects Behavioral and Neural Signatures of Model-Based Control during Sequential Decision Making.” Proceedings of the National Academy of Sciences of the United States of America 112 (5): 1595–1600.

Dezfouli, Amir, Nura W. Lingawi, and Bernard W. Balleine. 2014. “Habits as Action Sequences: Hierarchical Action Control and Changes in Outcome Value.” *Philosophical Transactions of the Royal Society of London. Series B*, Biological Sciences 369 (1655): 20130482.

Dolan, Ray J., and Peter Dayan. 2013. “Goals and Habits in the Brain.” Neuron 80 (2): 312–25.

Gera, Rani, Maya Bar Or, Ido Tavor, Dana Roll, Jeffrey Cockburn, Segev Barak, Elizabeth Tricomi, John P. O’Doherty, and Tom Schonberg. 2023. “Characterizing Habit Learning in the Human Brain at the Individual and Group Levels: A Multi-Modal MRI Study.” NeuroImage 272 (May): 120002.

Gera, Rani, Segev Barak, and Tom Schonberg. 2024. “A Novel Free-Operant Framework Enables Experimental Habit Induction in Humans.” Behavior Research Methods 56 (4): 3937–58.

Gillan, Claire M., Michal Kosinski, Robert Whelan, Elizabeth A. Phelps, and Nathaniel D. Daw. 2016. “Characterizing a Psychiatric Symptom Dimension Related to Deficits in Goal-Directed Control.” ELife 5 (March). 10.7554/eLife.11305.

Glück, Valentina M., Katharina Zwosta, Uta Wolfensteller, Hannes Ruge, and Andre Pittig. 2021. “Costly Habitual Avoidance Is Reduced by Concurrent Goal-Directed Approach in a Modified Devaluation Paradigm.” Behaviour Research and Therapy 146 (November): 103964.

Guida, Pasqualina, Mario Michiels, Peter Redgrave, David Luque, and Ignacio Obeso. 2022. “An FMRI Meta-Analysis of the Role of the Striatum in Everyday-Life vs Laboratory-Developed Habits.” Neuroscience and Biobehavioral Reviews 141 (104826): 104826.

Hardwick, Robert M., Alexander D. Forrence, John W. Krakauer, and Adrian M. Haith. 2019. “Time-Dependent Competition between Goal-Directed and Habitual Response Preparation.” Nature Human Behaviour 3 (12): 1252–62.

Hardwick, Robert M., Claudia Rottschy, R. Chris Miall, and Simon B. Eickhoff. 2013. “A Quantitative Meta-Analysis and Review of Motor Learning in the Human Brain.” NeuroImage 67 (February): 283–97.

Hogarth, L., C. Lam-Cassettari, and H. Pacitti. 2019. “Intact Goal-directed Control in Treatment-seeking Drug Users Indexed by Outcome-devaluation and Pavlovian to Instrumental Transfer: Critique of Habit Theory.” European Journal Of. https://onlinelibrary.wiley.com/doi/abs/10.1111/ejn.13961.

Luque, David, Sara Molinero, Poppy Watson, Francisco J. López, and Mike E. Le Pelley. 2020. “Measuring Habit Formation through Goal-Directed Response Switching.” Journal of Experimental Psychology. General 149 (8): 1449–59.

Meier, Jacqueline Katharina, Bernhard P. Staresina, and Lars Schwabe. 2022. “Stress Diminishes Outcome but Enhances Response Representations during Instrumental Learning.” ELife 11 (July). 10.7554/eLife.67517.

Michiels, Mario, Vincent Man, David Luque, and Ignacio Obeso. 2025. “The Neural Basis of Habit Formation Measured in Goal-Directed Response Switching.” BioRxiv. 10.1101/2025.03.13.643040.

Nebe, Stephan, André Kretzschmar, Maike C. Brandt, and Philippe N. Tobler. 2024. “Characterizing Human Habits in the Lab.” *Collabra*. Psychology 10 (1): 92949.

Paparella, Giulia, Martina De Riggi, Antonio Cannavacciuolo, Davide Costa, Daniele Birreci, Massimiliano Passaretti, Luca Angelini, et al. 2024. “Interhemispheric Imbalance and Bradykinesia Features in Parkinson’s Disease.” Brain Communications 6 (1): fcae020.

Perez, Omar D., and Anthony Dickinson. 2020. “A Theory of Actions and Habits: The Interaction of Rate Correlation and Contiguity Systems in Free-Operant Behavior.” Psychological Review 127 (6): 945–71.

Pérez, Omar D., and Fabian A. Soto. 2020. “Evidence for a Dissociation between Causal Beliefs and Instrumental Actions.” Quarterly Journal of Experimental Psychology (2006) 73 (4): 495–503.

Pool, Eva R., Rani Gera, Aniek Fransen, Omar D. Perez, Anna Cremer, Mladena Aleksic, Sandy Tanwisuth, et al. 2022. “Determining the Effects of Training Duration on the Behavioral Expression of Habitual Control in Humans: A Multilaboratory Investigation.” Learning & Memory 29 (1): 16–28.

Redgrave, Peter, Manuel Rodriguez, Yoland Smith, Maria C. Rodriguez-Oroz, Stephane Lehericy, Hagai Bergman, Yves Agid, Mahlon R. DeLong, and Jose A. Obeso. 2010. “Goal-Directed and Habitual Control in the Basal Ganglia: Implications for Parkinson’s Disease.” Nature Reviews. Neuroscience 11 (11): 760–72.

Reed, Phil. 2001. “Human Schedule Performance with Hypothetical Monetary Reinforcement.” European Journal of Behavior Analysis 2 (2): 225–34.

Rushworth, Matthew F. S., Maryann P. Noonan, Erie D. Boorman, Mark E. Walton, and Timothy E. Behrens. 2011. “Frontal Cortex and Reward-Guided Learning and Decision-Making.” Neuron 70 (6): 1054–69.

Seger, Carol A., and Brian J. Spiering. 2011. “A Critical Review of Habit Learning and the Basal Ganglia.” Frontiers in Systems Neuroscience 5 (August): 66.

Sjoerds, Zsuzsika, Anja Dietrich, Lorenz Deserno, Sanne de Wit, Arno Villringer, Hans-Jochen Heinze, Florian Schlagenhauf, and Annette Horstmann. 2016. “Slips of Action and Sequential Decisions: A Cross-Validation Study of Tasks Assessing Habitual and Goal-Directed Action Control.” Frontiers in Behavioral Neuroscience 10 (December): 234.

Smith, Kyle S., and Ann M. Graybiel. 2013. “A Dual Operator View of Habitual Behavior Reflecting Cortical and Striatal Dynamics.” Neuron 79 (2): 361–74.

Smith, Kyle S., and Ann M. Graybiel. 2016. “Habit Formation Coincides with Shifts in Reinforcement Representations in the Sensorimotor Striatum.” Journal of Neurophysiology 115 (3): 1487–98.

Watson, P., T. E. Gladwin, A. A. C. Verhoeven, and S. de Wit. 2023. “Investigating Habits in Humans with a Symmetrical Outcome-Revaluation Task.” Behavior Research Methods 55 (5): 2687–2705.

Watson, Poppy. 2024. “Defining and Measuring Habits across Different Fields of Research.” In Habits, 3–22. Cham: Springer International Publishing.

Watson, Poppy, Claire O’Callaghan, Iain Perkes, Laura Bradfield, and Karly Turner. 2022. “Making Habits Measurable beyond What They Are Not: A Focus on Associative Dual-Process Models.” Neuroscience & Biobehavioral Reviews 142.

Weintraub, Daniel, Juergen Koester, Marc N. Potenza, Andrew D. Siderowf, Mark Stacy, Valerie Voon, Jacqueline Whetteckey, Glen R. Wunderlich, and Anthony E. Lang. 2010. “Impulse Control Disorders in Parkinson Disease: A Cross-Sectional Study of 3090 Patients.” Archives of Neurology 67 (5): 589–95.

Wit, Sanne de, Merel Kindt, Sarah L. Knot, Aukje A. C. Verhoeven, Trevor W. Robbins, Julia Gasull-Camos, Michael Evans, Hira Mirza, and Claire M. Gillan. 2018. “Shifting the Balance between Goals and Habits: Five Failures in Experimental Habit Induction.” Journal of Experimental Psychology. General 147 (7): 1043–65.

